# Dynamic endothelial ephrin-B2 and EphB4 expression characterizes hyaloid vessel regression

**DOI:** 10.64898/2026.10.17.758877

**Authors:** Zouine Kawtar, Sahrout Walid Miguel, Megne Ariane Vanessa, Oubaha Malika

## Abstract

Arteriovenous endothelial identity coordinates vascular growth and patterning, but whether it is remodeled during developmental vessel regression remains unknown. The mouse hyaloid vasculature is a transient ocular network considered predominantly arterial; its timely regression is required for normal vision, yet the identity of its regressing endothelium has remained largely unexplored. Here, we show that postnatal regression is accompanied by a shift from ephrin-B2-enriched to EphB4-enriched endothelium. Residual ephrin-B2-positive endothelial cells increasingly displayed cleaved caspase-3 labeling, whereas Ki67 positivity increased among EphB4-positive cells. Endothelial-specific Notch1 deletion, which causes persistent hyaloid vessels, accentuated this ephrin-B2-low/EphB4-high profile while reducing apoptotic labeling within residual ephrin-B2-positive cells and increasing proliferation within the EphB4-positive population. Mosaic recombination further showed that recombined endothelial cells preferentially expressed EphB4, whereas neighboring unrecombined cells more frequently retained ephrin-B2, supporting a cell-autonomous contribution of Notch1. Thus, arteriovenous-associated endothelial features remain dynamic during developmental vessel regression, linking endothelial identity to the cellular behaviors accompanying hyaloid remodeling.

**One Sentence Summary:** Dynamic shifts in endothelial ephrin-B2 and EphB4 expression accompany hyaloid vessel regression, with ephrin-B2⁺ ECs preferentially undergoing apoptosis and EphB4⁺ ECs exhibiting increased proliferation.

## INTRODUCTION

The formation of a functional vascular network requires endothelial cells (ECs) to acquire distinct arterial and venous identities. These identities are initiated by genetically encoded programs before the onset of circulation and are subsequently refined by local molecular and haemodynamic cues[1–4]. Although arterial and venous ECs were historically considered to represent stable lineages, their identities remain plastic during vascular development and remodeling[5]. Venous ECs can, for example, be developmentally reprogrammed to form coronary arteries, whereas inappropriate perturbation of arteriovenous identity in established vascular beds can contribute to vascular malformations[6]. Thus, the maintenance and context-dependent adaptation of endothelial arteriovenous identity are essential for normal vascular development and homeostasis[7].

Notch signaling is a central regulator of arterial specification[8]. Vascular endothelial growth factor (VEGF) induces components of the Notch pathway, whereas activated endothelial Notch promotes arterial differentiation and suppresses venous characteristics[2, 9–11]. Conversely, loss of Notch signaling reduces the expression of arterial markers, including ephrin-B2, and disrupts arteriovenous patterning[2, 9, 12, 13]. Ephrin-B2 and its receptor EphB4 are enriched in arterial and venous ECs, respectively, from early embryonic development[6, 14–16]. However, these molecules are not simply markers of fixed endothelial lineages. Eph-ephrin signaling regulates cell adhesion, repulsion, migration and segregation[17], thereby contributing to the positioning of arterial- and venous-fated ECs within developing vascular networks[2, 14]. Moreover, ephrin-B2 and EphB4 are not expressed exclusively in mature arterial and venous domains: both can be detected in angiogenic ECs, and their relative expression influences endothelial plasticity and arteriovenous specification. Recent work demonstrated that the reciprocal balance between ephrin-B2 and EphB4 couples EC sorting to arterial specification through interactions with VEGF, Notch and flow-responsive programs[18]. EphB4 restrains arterial specification, whereas increased ephrin-B2 activity promotes the acquisition and positioning of arterial ECs. Disruption of this balance leads to abnormal arteriovenous patterning, emphasizing that the relative expression and activities of ephrin-B2 and EphB4 contribute to endothelial identity[18].

Whether endothelial arteriovenous identity is similarly remodeled during developmental vessel regression remains unknown. The hyaloid vasculature is a transient fetal network that supports the developing lens and inner eye before undergoing postnatal regression[19, 20]. Its arteriovenous organization differs among vertebrate species. Although the mouse hyaloid vasculature has traditionally been described as an arterial network, this classification is based predominantly on its anatomical origin from the hyaloid artery. Whether its endothelial cells maintain a uniform arterial identity during regression, and whether comparable arteriovenous programs operate in the human fetal hyaloid vasculature, remain unknown[21–23]. This represents an important gap because hyaloid vessel regression is an active remodeling process involving coordinated changes in endothelial proliferation, positioning and interactions with mural and immune cells[24, 25]. Failure of this process causes persistent fetal vasculature (PFV), a congenital developmental anomaly in which remnants of the fetal hyaloid system persist and can cause severe visual impairment[26–30].

We recently showed that endothelial Notch1 is required for normal hyaloid vessel regression. Endothelial-specific deletion of Notch1 results in excessive EC proliferation, impaired endothelial disengagement and defective multicellular remodeling, leading to persistent hyaloid vessels[31]. These findings identified Notch1 as a key regulator of endothelial apoptosis and proliferation during hyaloid vessel regression. However, whether this process is accompanied by changes in endothelial arteriovenous identity remains unknown. This question is particularly relevant given the role of endothelial specification in vascular remodeling. Notch1 promotes arterial specification and ephrin-B2 expression, whereas EphB4 is associated with venous endothelial identity and proliferative states[2, 32]. We therefore examined the temporal changes in endothelial ephrin-B2 and EphB4 expression during physiological hyaloid regression and determined how their relative distribution is affected by endothelial Notch1 loss.

Here, we show that mouse hyaloid vessels undergo progressive changes in endothelial arteriovenous marker expression during postnatal regression. Ephrin-B2 expression and the proportion of ephrin-B2⁺ ECs decline, whereas EphB4 expression and the EphB4⁺ endothelial population increase. These populations exhibit distinct patterns of apoptosis and proliferation, with ephrin-B2⁺ ECs showing greater apoptotic labeling and EphB4⁺ ECs accounting for most proliferating ECs. Endothelial-specific Notch1 deletion further reduces ephrin-B2 expression and increases the proportion of EphB4⁺ ECs. Mosaic analysis reveals preferential EphB4 expression in Notch1-deficient ECs, consistent with a cell-autonomous contribution of Notch1 to endothelial marker regulation. Together, these findings identify Notch1-dependent changes in endothelial ephrin-B2/EphB4 expression that accompany hyaloid vessel regression and are altered in persistent vessels following Notch1 loss. Whether these changes contribute directly to vessel persistence remains to be established.

## RESULTS

### Hyaloid vessel regression is accompanied by a shift in arteriovenous-associated marker expression

To determine whether arteriovenous-associated expression changes during hyaloid vessel regression, we analyzed isolated mouse hyaloid vessels at embryonic day (E)18.5 and postnatal days (P)0, P4 and P8, spanning the onset and progression of postnatal regression (Fig. 1A). Quantitative RT-PCR showed that *Efnb2* expression decreased sharply after E18.5 and remained reduced throughout the postnatal period. By contrast, *Ephb4* expression was not significantly altered at P0 or P4 but increased markedly by P8 (Fig. 1B).

**Figure 1.**
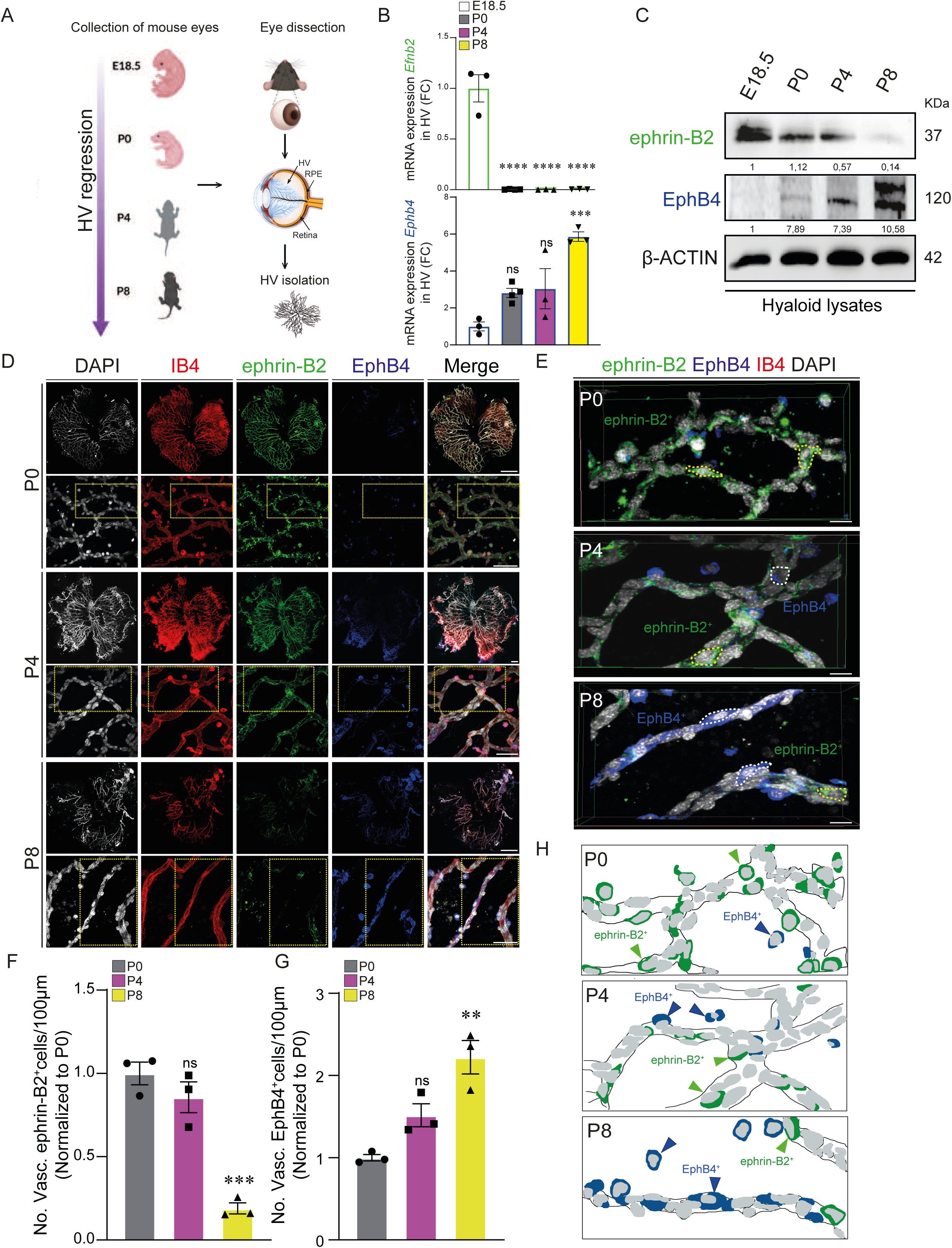
Hyaloid vessel regression is accompanied by decreased ephrin-B2 and increased EphB4 expression. (A) Experimental workflow for hyaloid vessel (HV) collection and isolation at embryonic day 18.5 (E18.5), and postnatal day 0 (P0), P4, and P8. (B) Quantitative reverse transcription polymerase chain reaction (qRT-PCR) of arteriovenous identity genes at E18.5, P0, P4 and P8 reveals a downregulation of arterial transcript *Efnb2* and progressive upregulation of venous gene *EphB4* as regression progresses (*n* = 3). (C) Western blot analysis of hyaloid cell lysates at P0, P4 and P8 showing decreased ephrin-B2 and increased EphB4 protein levels at later stages (*n*=2-3). Densitometric quantification of each protein is shown relative to the loading control, β-ACTIN. (D) Representative confocal micrographs and (E) 3D reconstructions of P0, P4, and P8 HV flatmounts labeled for ephrin-B2 (green), EphB4 (blue), IB4 (red), and DAPI (white) (*n*=5-6). (F-G) Quantification of vascular ephrin-B2⁺ (F) and EphB4⁺ (G) cells per 100 µm of vessel length in HV at P0, P4, and P8 (*n* = 3). (H) Model summarizing the dynamic distribution of ephrin-B2⁺ and EphB4⁺ cells during HV regression. Results are expressed as fold change (FC) relative to E18.5 (B) and as cell numbers (No) per 100 µm of vessel length normalized to P0 (F, G). Data are shown as means ± SEM. Represented *p* values are **≤ 0.01, ***≤ 0.001, ****≤0.0001 from ordinary one-way ANOVA test with Dunnett’s multiple comparison (B, F, G). Non-significant (ns). Scale bars, 200 μm (D) and 20 μm [for higher magnification images in (D, E)].

Analysis of protein abundance revealed related but distinct temporal profiles. Ephrin-B2 protein remained abundant from E18.5 to P4 but was markedly reduced at P8. Conversely, EphB4 protein was weakly detected at E18.5, increased by P0 and remained elevated throughout postnatal regression (Fig. 1C). Thus, although transcript and protein abundance displayed different kinetics, both analyses identified a late-regression profile characterized by reduced ephrin-B2 and increased EphB4 expression.

To determine whether these changes extended beyond ephrin-B2 and EphB4, we analyzed additional arteriovenous-associated genes. The arterial-associated genes *Notch1* and *Dll4* showed transient increases during early postnatal development followed by reduced expression at P8, whereas *Gja5* decreased after E18.5. Conversely, expression of the venous-associated genes *Aplnr* and *Nr2f2* increased during postnatal regression, while *Tek* was strongly upregulated at P8 (Fig. S1A). These marker-specific kinetics indicate that hyaloid vessel regression is not accompanied by a synchronous binary switch of the entire arteriovenous program. Instead, late regression is characterized by a broader reduction in arterial-associated expression and enrichment of venous-associated transcripts.

We next examined the spatial distribution of ephrin-B2 and EphB4 in hyaloid whole mounts. In the P8 retina, ephrin-B2 was enriched in arteries, whereas EphB4 was predominantly associated with veins, providing an anatomical control for their expected vascular distribution (Fig. S1B). In the hyaloid vasculature, ephrin-B2-positive cells were broadly associated with the IB4⁺ vascular network at P0, whereas EphB4-positive cells were infrequent. Both markers were detected within the remodeling network at P4, with a relative predominance of ephrin-B2. By P8, vascular-associated ephrin-B2 labeling was markedly reduced, whereas EphB4-positive cells were more abundant along the remaining vessels (Fig. 1D, E).

Quantification confirmed that the abundance of vascular-associated ephrin-B2-positive cells was unchanged between P0 and P4 but markedly reduced at P8. Conversely, EphB4-positive cell abundance did not differ significantly between P0 and P4 but increased approximately twofold by P8 (Fig. 1F, G). These changes are summarized schematically in Fig. 1H. Together, these findings show that late-stage hyaloid vessel regression is associated with a marker-specific shift from an ephrin-B2-enriched toward an EphB4-enriched vascular profile.

### Endothelial ephrin-B2 and EphB4 exhibit distinct expression dynamics during hyaloid vessel regression

Because ephrin-B2 and EphB4 are not restricted to the vascular endothelium, we next determined whether the tissue-level changes identified during hyaloid vessel regression were reflected within ECs. Hyaloid vessels were co-immunostained for ephrin-B2 or EphB4 together with the endothelial nuclear marker ERG at P0, P4 and P8 (Fig. 2).

**Figure 2.**
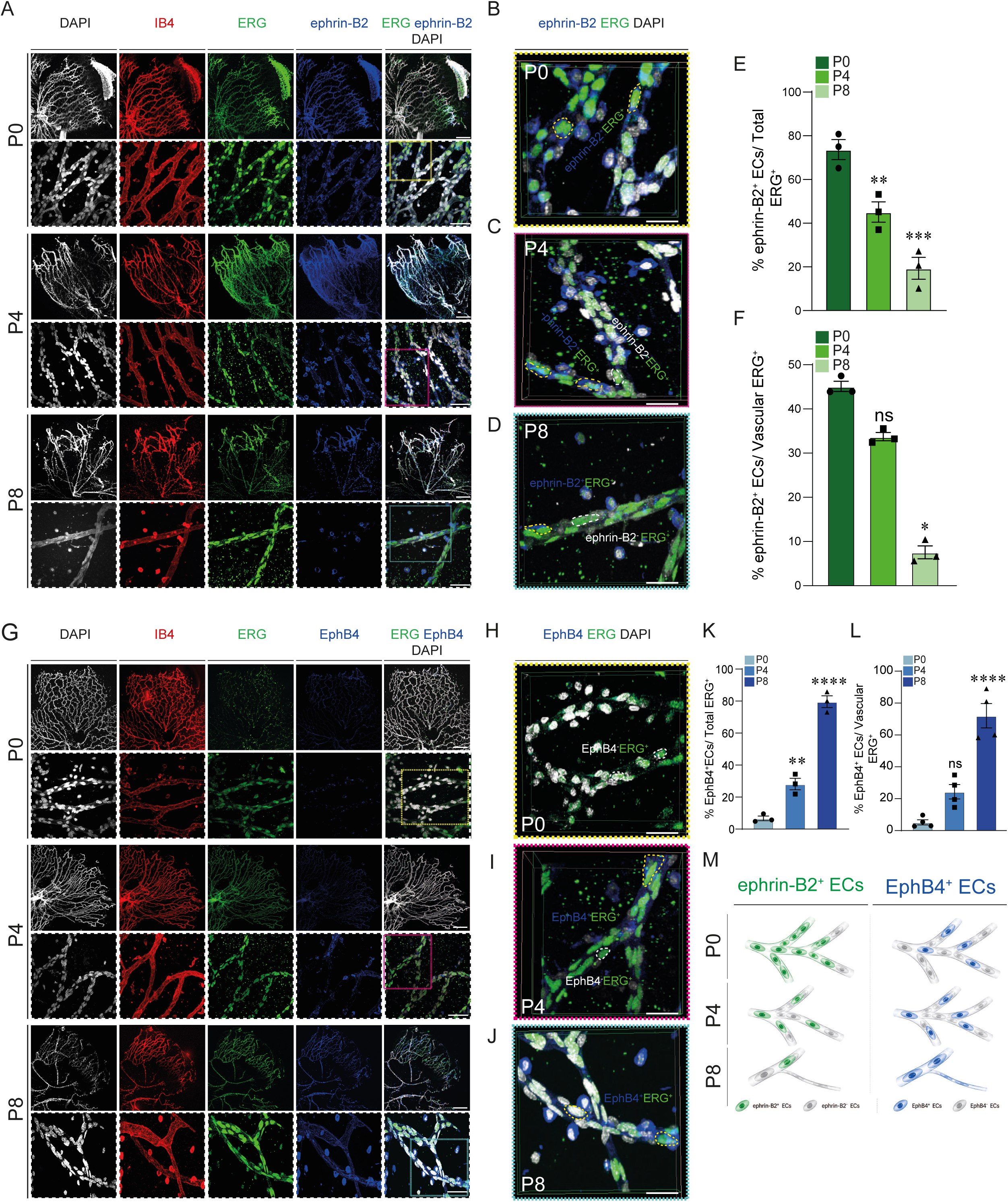
Endothelial ephrin-B2 and EphB4 expression changes during hyaloid vessel regression. (A) Representative confocal images of whole-mount HV immunostained for IB4 (red), ERG1/2/3 (green), and ephrin-B2 (blue) at P0, P4 and P8. Merged images are shown with DAPI (white). Boxed regions are shown at higher magnification below. (B-D) Representative 3D reconstructions of confocal z-stacks illustrating ephrin-B2 expression in ERG1/2/3⁺ endothelial cells (ECs) within the hyaloid vasculature at P0 (B), P4(C), and P8(D). (E) Quantification of ephrin-B2^+^ EC relative to total number of ERG1/2/3^+^ cells in HV at P0, P4, and P8 (*n* =3). (F) Quantification of ephrin-B2^+^ ECs expressed relative to the total number of vascular ERG1/2/3^+^ cells at P0, P4, and P8 (*n* =3). (G) Representative confocal micrographs of IB4 (red), ERG1/2/3 (green), EphB4 (blue), and DAPI (white) at P0, P4 and P8 reveal a progressive increase in ERG1/2/3⁺EphB4⁺ cells during HV regression. (H-J) 3D reconstitution confirmed the localization of EphB4 within ERG1/2/3⁺ cells at P0 (H), P4(I), and P8(J). (K) Quantification of EphB4^+^ ECs among total ERG1/2/3⁺ cells at P0, P4, and P8 (*n* = 3). (L) Quantification of EphB4⁺ ECs among vascular ERG1/2/3⁺ cells at P0, P4, and P8 (*n*=3). (M) Model summarizing the temporal changes in ephrin-B2^+^, and EphB4^+^ cells at P0, P4 and P8. Ephrin-B2 and EphB4 profiles were quantified from separate immunostainings and therefore do not represent mutually exclusive endothelial populations. Data are shown as means ± SEM. Represented *p* values are *≤ 0.05, **≤ 0.01, ***≤ 0.001, ****≤0.0001 from ordinary one-way ANOVA with Dunnett’s multiple comparisons test (E, K, L), Kruskal-Wallis test with Dunn’s multiple comparisons test (F). Non-significant (ns). Scale bars, 500 μm (A, G) and 50 μm [for higher magnification images in (A-D, G-J)].

At P0, ephrin-B2 was broadly detected in ERG⁺ ECs throughout the hyaloid vascular network (Fig. 2A, B). As regression proceeded, the abundance of ephrin-B2⁺ERG⁺ ECs decreased, as illustrated by three-dimensional (3D) reconstructions at P0, P4 and P8 (Fig. 2B-D). Quantification showed that the proportion of ephrin-B2⁺ ECs among total ERG⁺ ECs significantly decreased at P4 and P8 relative to P0 (Fig. 2E). Within the vascular ERG⁺ population, ephrin-B2 positivity declined from approximately 45% at P0 to 33% at P4 and 6% at P8, reaching statistical significance at P8 (Fig. 2F). Thus, ephrin-B2⁺ ECs are progressively depleted from the hyaloid vasculature, with the greatest reduction occurring during late regression.

In contrast, EphB4⁺ERG⁺ ECs were infrequent at P0 but became increasingly prominent as regression progressed (Fig. 2G-J). The proportion of EphB4⁺ cells among all ERG⁺ ECs were significantly increased at P4 and P8 compared with P0 (Fig. 2K). Within the vascular ERG⁺ population, EphB4 positivity increased from approximately 5% at P0 to 26% at P4 and 78% at P8, reaching statistical significance at P8 (Fig. 2L). Thus, EphB4⁺ ECs become progressively enriched within the hyaloid vasculature, with the greatest increase occurring during late regression.

We next assessed ephrin-B2 and EphB4 expression in ERG⁺ cells located outside IB4⁺ vascular structures. The proportion of extravascular ERG⁺ cells expressing ephrin-B2 did not change significantly between P0 and P8 (Fig. S2A). By contrast, EphB4 expression within the extravascular ERG⁺ population increased significantly at P4 and remained elevated at P8 (Fig. S2B). Thus, EphB4 enrichment during regression was detected in both vessel-associated and extravascular ERG⁺ endothelial populations (Fig. S2B).

Because ephrin-B2 and EphB4 were also detected in ERG⁻ cells, we examined their expression in F4/80⁺ hyaloid-associated macrophages. Ephrin-B2 was detected in approximately 90% of F4/80⁺ cells at both P0 and P8, with no significant temporal change (Fig. S2C, D). In contrast, the proportion of EphB4-positive macrophages increased from approximately 30% at P0 to more than 90% at P8 (Fig. S2E, F). These findings confirm that ephrin-B2 and EphB4 are not endothelial-specific and underscore the importance of ERG co-localization for defining their expression within ECs.

Together, these findings reveal a progressive shift in the arteriovenous marker profile of the hyaloid endothelium, from an early predominance of ephrin-B2⁺ ECs to an enrichment of EphB4⁺ ECs during late regression, highlighting dynamic changes in endothelial marker expression throughout vascular regression.

### Ephrin-B2⁺ and EphB4⁺ endothelial populations display distinct apoptotic and proliferative profiles during hyaloid vessel regression

Having established that the regressing hyaloid endothelium shifts from an ephrin-B2-enriched to an EphB4-enriched profile, we next asked whether these marker-defined endothelial populations were associated with distinct cellular behaviors. We examined apoptosis and proliferation at P0, P4 and P8 using cleaved caspase-3 (C-Casp3) and Ki67, respectively.

To assess apoptosis within the ephrin-B2-positive endothelial population, hyaloid vessels were co-immunostained for ephrin-B2, CD31 and C-Casp3 (Fig. 3A). Three-dimensional reconstructions confirmed the localization of C-Casp3 within ephrin-B2⁺CD31⁺ cells within the vessel wall (Fig. 3B). The number of apoptotic ephrin-B2⁺ ECs per 1 00 µm of vessel length significantly increased at P4 and P8 compared with P0 (Fig. 3C). Within the ephrin-B2⁺ endothelial population, the proportion of C-Casp3-positive cells increased from approximately 5% at P0 to 11% at P4 and 84% at P8 (Fig. 3D). Thus, the late reduction in ephrin-B2-positive ECs coincided with a marked enrichment of apoptotic cells within this population.

**Figure 3.**
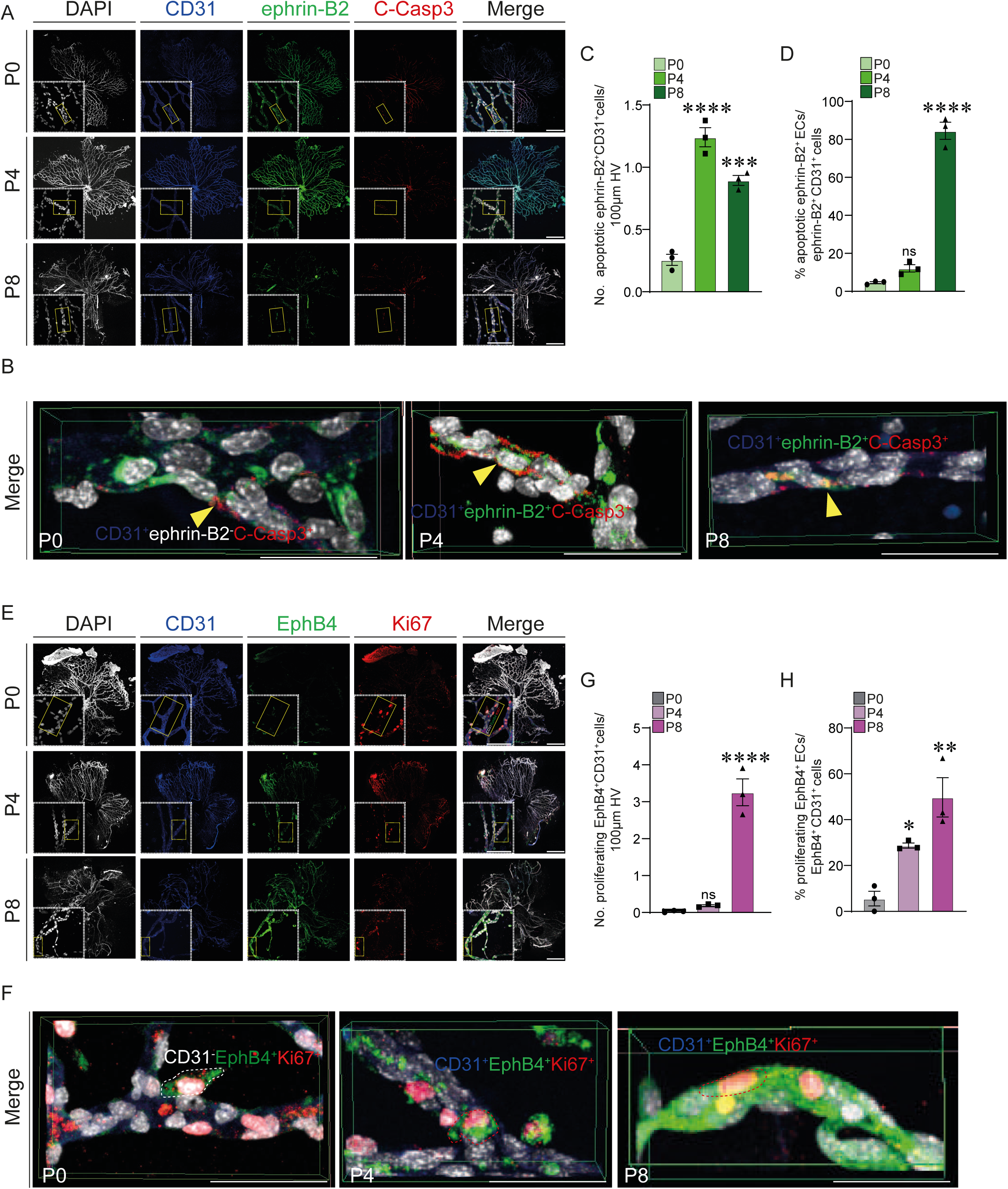
Differential proliferative and apoptotic activities of EphB4⁺ and ephrin-B2⁺ endothelial cells during hyaloid vessel regression. (A) Confocal images of CD31 (blue), ephrin-B2 (green), C-Casp3 (red), and DAPI (white) co-immunostaining at P0, P4 and P8 reveal apoptotic ephrin-B2⁺ ECs during HV regression. (B) 3D reconstitution showing apoptotic ephrinB2^+^ ECs (*n* =4). (C) Quantification of apoptotic ephrin-B2⁺ ECs (C-Casp3⁺ephrin-B2⁺CD31⁺) per 100 µm of vessel length at P0, P4, and P8. (D) Percentage of C-Casp3⁺ephrin-B2⁺CD31⁺ ECs among ephrin-B2⁺CD31⁺ ECs at P0, P4, and P8 (*n* = 3). (E) Representative confocal micrographs of whole-mount hyaloid vessels immunostained for CD31 (blue), EphB4(green), Ki67(red) and DAPI (white) at P0, P4 and P8. (F) Representative 3D reconstitution showing proliferating EphB4^+^ ECs within the hyaloid vasculature (*n* =4). (G) Quantification of proliferating EphB4⁺ ECs (Ki67⁺EphB4⁺CD31⁺) per 100 µm of vessel length at P0, P4, and P8. (H) Percentage of Ki67⁺EphB4⁺CD31⁺ ECs among EphB4⁺CD31⁺ ECs at P0, P4, and P8 (*n* = 3). Data are shown as means ± SEM. Represented *p*-values are *≤ 0.05, **≤ 0.01, and ****≤0.0001 from ordinary one-way ANOVA with Dunnett’s multiple comparisons test. Non-significant (NS). Scale bars, 500µm (A, E) and 50µm [for higher-magnification images in (A, B, E, F)].

We next determined whether the expansion of EphB4-positive ECs was associated with endothelial proliferation. Hyaloid vessels were co-immunostained for EphB4, CD31 and Ki67 (Fig. 3E), and 3D reconstruction confirmed the presence of Ki67 within EphB4⁺CD31⁺ ECs (Fig. 3F). The number of proliferating EphB4⁺ ECs per 100 µm of vessel length remained low and comparable at P0 and P4 but increased markedly at P8 (Fig. 3G). Similarly, the proportion of Ki67⁺ cells within the EphB4⁺ endothelial population progressively increased from approximately 5% at P0 to 28% at P4 and 50% at P8 (Fig. 3H). Notably, this increase contrasted with the overall decline in endothelial proliferation, indicating that EphB4⁺ ECs exhibited enhanced proliferative activity during the later stages of hyaloid vessel regression. Together, these findings reveal distinct temporal associations between arteriovenous-associated endothelial markers and cellular behavior during hyaloid vessel regression. Apoptotic cells become enriched within the declining ephrin-B2-positive population, whereas proliferation increases within the expanding EphB4-positive population. These complementary associations accompany, but do not by themselves establish the cause of, the population-level shift from an ephrin-B2-enriched to an EphB4-enriched endothelial profile.

### Endothelial Notch1 loss alters arteriovenous marker expression and leads to hyaloid vessels persistence

Notch signaling promotes arterial endothelial specification, and we previously showed that endothelial-specific deletion of *Notch1* impairs hyaloid vessel regression. We therefore asked whether endothelial Notch1 loss alters the expression of arteriovenous-associated markers in persistent hyaloid vessels. *Notch1* was deleted postnatally using *Cdh5-CreERT2; Notch1^fl/fl^*mice, with tamoxifen administered from P1 to P3 and hyaloid vessels analyzed at P8 (Fig. 4A).

**Figure 4.**
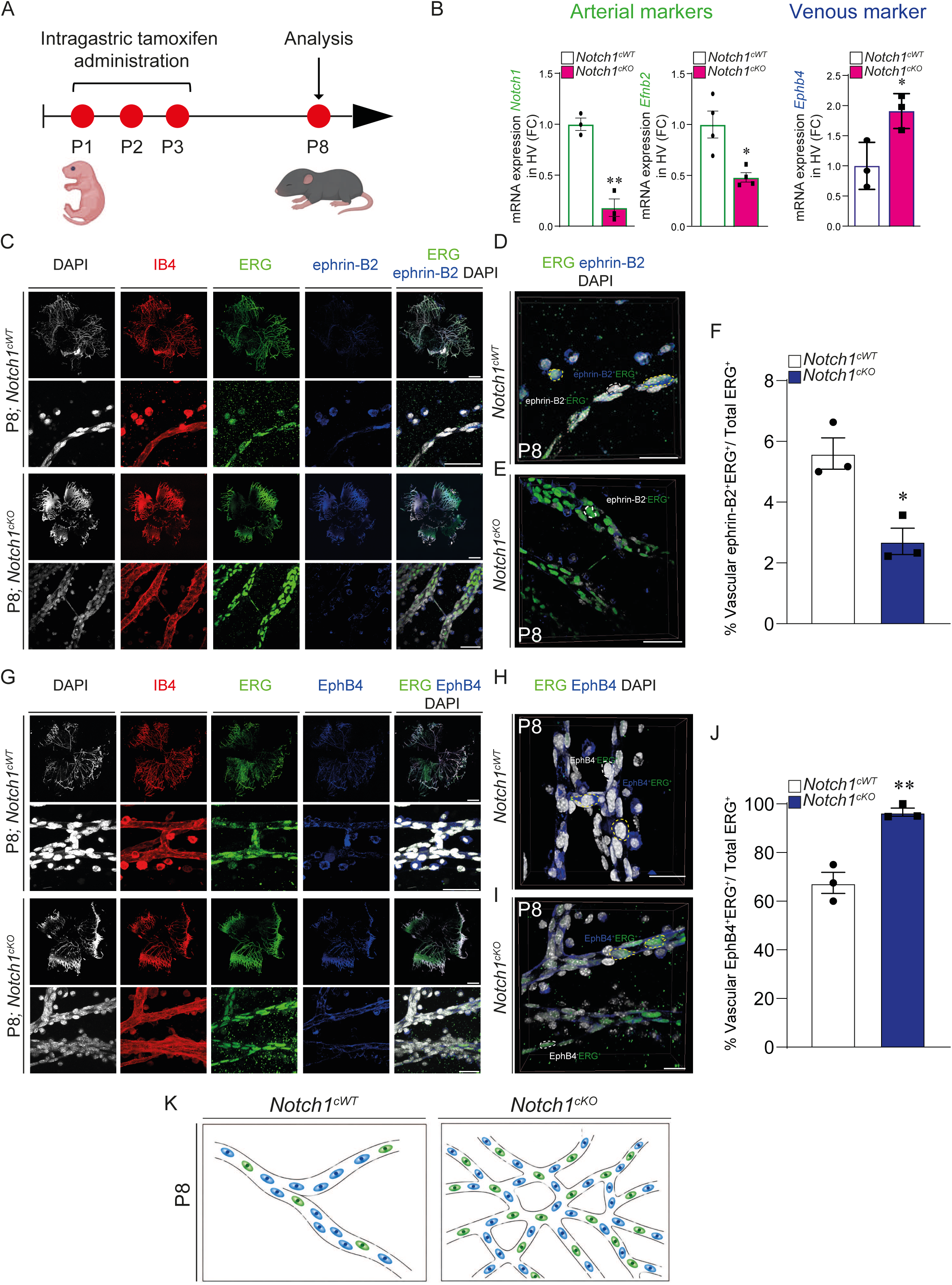
Impaired hyaloid vessel regression is associated with an arteriovenous identity imbalance. (A) Experimental design illustrating tamoxifen administration (15 µg/pup, 30 µL) to induce endothelial-specific conditional deletion of *Notch1*, followed by HV analysis at P8. (B) Quantitative RT-PCR analysis of P8 HV from *Notch1^cWT^ and Notch1^cKO^* mice showing reduced expression of the arterial markers *Notch1* and *Efnb2*, and increased expression of the venous marker *Ephb4* following endothelial *Notch1* deletion (*n* =3). (C) Representative confocal micrographs and (D, E) 3D reconstructions from *Notch1^cWT^* and *Notch1^cKO^*mice labeled for IB4 (red), ERG (green), ephrin-B2 (blue), and DAPI (white), illustrating ephrin-B2^+^ERG1/2/3^+^ (yellow circle) and ephrin-B2^−^ERG1/2/3^+^ (white circle) endothelial populations (*n* =3). (F) Quantification of the proportion of vascular ephrin-B2⁺ ECs among total ERG1/2/3⁺ cells in P8 HV from *Notch1^cW^*^T^ and *Notch1^cKO^* mice (*n* = 3). (G) Representative confocal images of P8 HV co-immunostained for IB4 (red), ERG (green), EphB4 (blue), and DAPI (white) in *Notch1^cWT^* and *Notch1^cKO^* mice. (H, I) Representative 3D reconstructions illustrating EphB4⁺ERG1/2/3⁺ cells (yellow circle), and EphB4⁻ERG1/2/3⁺ (white circle) endothelial populations (*n* = 3). (J) Quantification of the proportion of vascular EphB4^+^ ECs among total vascular ERG1/2/3⁺ ECs in P8 HV from *Notch1^cWT^* and *Notch1^cKO^* mice (*n* = 3). (K) Schematic summary illustrating the relative changes in endothelial ephrin-B2 and EphB4 expression associated with endothelial *Notch1* deletion. Ephrin-B2 and EphB4 profiles were quantified from separate immunostainings and therefore do not represent mutually exclusive endothelial populations Data are presented as means ± SEM. Represented *p*-values are *≤ 0.05 and **≤ 0.01 from two-tailed parametric unpaired t-test. Non-significant (NS). Scale bars, 500 μm (C, G) and 50µm [for higher-magnification images in (C-E, G-I)].

Quantitative RT-PCR of isolated hyaloid vessels confirmed a marked reduction in *Notch1* expression in *Notch1^cKO^* mice. This was accompanied by decreased expression of the arterial-associated gene *Efnb2* and increased expression of the venous-associated gene *Ephb4* (Fig. 4B). Consistent with a broader alteration of the arteriovenous expression profile, *Aplnr* and *Tek* expression increased, whereas expression of the arterial-associated gene *Dll4* decreased in *Notch1*^cKO^ hyaloid vessels (Fig. S3A).

To determine whether the changes in ephrin-B2 and EphB4 occurred within the endothelium, P8 hyaloid vessels were co-immunostained for ephrin-B2, ERG and IB4. Ephrin-B2 was detected in vascular ERG⁺ endothelial cells in both genotypes, but the proportion of ephrin-B2⁺ cells among vascular ERG⁺ endothelial cells was significantly reduced in *Notch1^cKO^* vessels (Fig. 4C-F).

Conversely, co-immunostaining for EphB4, ERG and IB4 revealed an increased representation of EphB4⁺ endothelial cells in persistent *Notch1^cKO^* vessels. 3D reconstruction confirmed the endothelial localization of EphB4, and quantification demonstrated a significant increase in the proportion of EphB4⁺ cells among vascular ERG⁺ endothelial cells (Fig. 4G-J). Whole-mount co-immunostaining further illustrated reduced ephrin-B2 and enhanced EphB4 labeling in *Notch1^cKO^* hyaloid vessels (Fig. S3B). The schematic in Fig. 4K summarizes these findings at P8, illustrating the altered endothelial arteriovenous identity associated with hyaloid vessel persistence in *Notch1^cKO^* mice.

Together, these findings show that endothelial Notch1 deficiency accentuates an ephrin-B2-low/EphB4-high endothelial profile in persistent P8 hyaloid vessels. Thus, Notch1 is required for normal regulation of arteriovenous-associated marker expression during hyaloid vessel remodeling. Whether this altered expression profile contributes directly to vascular persistence cannot be determined from these experiments alone.

### Endothelial Notch1 loss alters apoptotic and proliferative profiles within ephrin-B2⁺ and EphB4⁺ endothelial populations

Physiological hyaloid vessel regression was associated with increased apoptotic labeling among ephrin-B2⁺ endothelial cells and increased proliferative activity among EphB4⁺ endothelial cells (Fig. 3). Because endothelial Notch1 deletion accentuated the ephrin-B2-low/EphB4-high profile but resulted in persistent hyaloid vessels, we asked whether the cellular behaviors associated with these endothelial populations were also altered.

To examine apoptosis within the residual ephrin-B2⁺ endothelial population, P8 hyaloid vessels from *Notch1^cWT^* and *Notch1^cKO^* mice were co-immunostained for ephrin-B2, C-Casp3 and CD31 (Fig. 5A). Three-dimensional reconstructions confirmed the localization of C-Casp3 signal within ephrin-B2⁺CD31⁺ endothelial cells (Fig. 5B). The proportion of C-Casp3-positive cells among ephrin-B2⁺ endothelial cells was significantly lower in *Notch1*^cKO^ vessels than in controls (Fig. 5C). Thus, the reduced abundance of ephrin-B2⁺ endothelial cells following Notch1 deletion was not accompanied by increased apoptotic labeling within the residual ephrin-B2⁺ population at P8.

**Figure 5.**
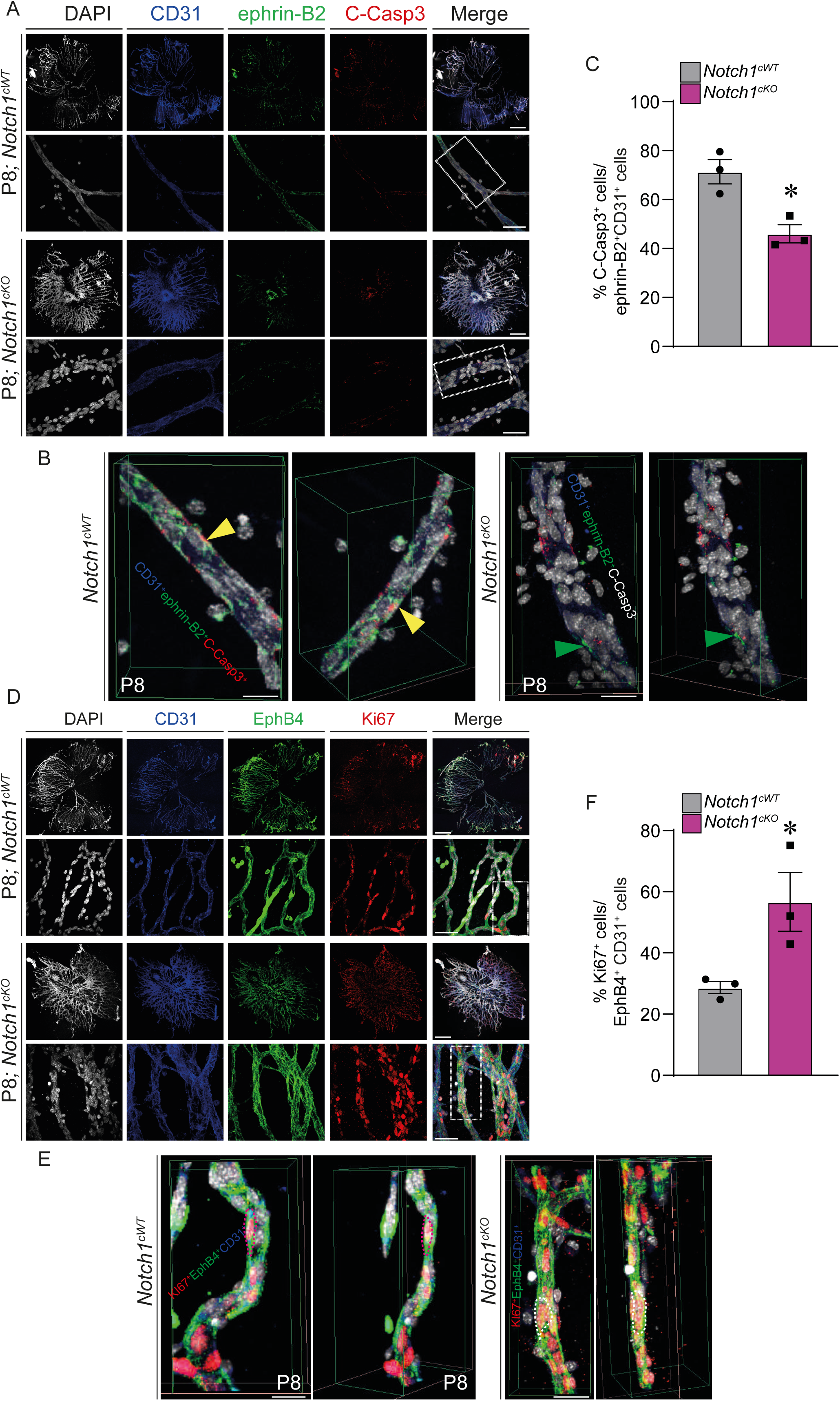
Endothelial Notch1 loss is associated with reduced apoptosis among ephrin-B2^+^ endothelial cells and increased proliferation among EphB4^+^ endothelial cells. (A) Representative confocal micrographs of P8 HV from *Notch1^cWT^*and *Notch1^cKO^* co-immunostained for CD31 (blue), ephrin-B2 (green), C-Casp3 (red) and DAPI (white) (*n* = 4). (B) 3D reconstructions showing apoptotic ephrin-B2⁺ ECs (C-Casp3⁺ephrin-B2⁺CD31⁺). (C) Quantification of C-Casp3⁺ cells among ephrin-B2⁺ CD31^+^ cells in P8 HV from *Notch1^cWT^* and *Notch1^cKO^*, normalized to *Notch1cWT* controls (*n* = 3). (D) Representative confocal micrographs of P8 HV from *Notch1^cWT^* and *Notch1^cKO^* co-immunostained for CD31 (blue), EphB4 (green), Ki67 (red) and DAPI (white) (*n*=3). (E) 3D reconstructions showing increased proliferation of EphB4⁺ ECs following endothelial *Notch1* deletion. (F) Quantification of Ki67⁺ cells among EphB4⁺ CD31+ in P8 HV from *Notch1^cWT^* and *Notch1^cKO^*mice (*n* = 3). Data are presented as means ± SEM. Represented *p*-values are *≤ 0.05 from two-tailed parametric unpaired t-test. Scale bars, 500 μm and 50 μm [for higher-magnification images in (A, B, D, E)].

We next assessed proliferation within the expanded EphB4⁺ endothelial population. Co-immunostaining for EphB4, Ki67 and CD31 identified proliferating EphB4⁺ endothelial cells in both genotypes (Fig. 5D). Three-dimensional reconstructions confirmed the localization of Ki67⁺ nuclei within EphB4⁺CD31⁺ vascular structures (Fig. 5E). The proportion of Ki67-positive cells among EphB4⁺ endothelial cells was significantly increased in *Notch1*^cKO^ vessels compared with controls (Fig. 5F). This finding is consistent with our previous demonstration of increased endothelial proliferation following endothelial Notch1 deletion[31].

Together, these findings show that endothelial Notch1 loss differentially affects apoptosis and proliferation across marker-defined endothelial populations in persistent hyaloid vessels. Residual ephrin-B2⁺ ECs exhibit reduced apoptotic labeling, whereas EphB4⁺ ECs show increased proliferative activity. These alterations accompany hyaloid vessel persistence, although their causal contribution remains to be established.

### Mosaic endothelial Notch1 deletion supports cell-autonomous regulation of ephrin-B2 and EphB4 expression

To investigate whether Notch1 regulates endothelial ephrin-B2 and EphB4 expression cell-autonomously, we induced mosaic recombination in *Rosa26-mT/mG; Cdh5-CreERT2; Notch1^fl/fl^* mice. Cre-mediated recombination switches membrane tdTomato (tdT) expression to GFP, allowing recombined and unrecombined endothelial cells to be compared within the same hyaloid vascular network (Fig. 6A).

**Figure 6.**
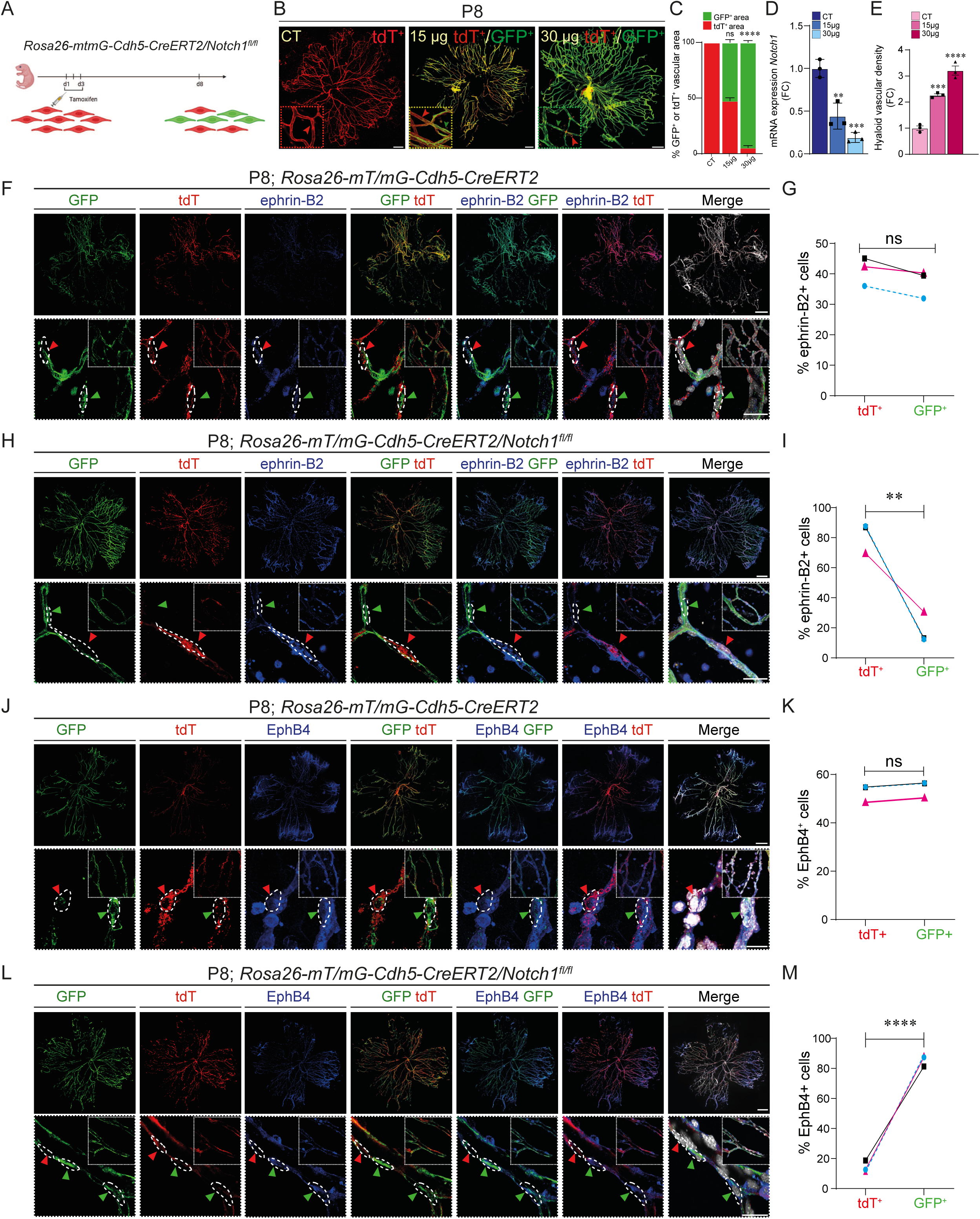
Mosaic endothelial Notch1 deletion is associated with altered ephrin-B2 and EphB4 expression. (A) Experimental design showing tamoxifen administration to induce mosaic endothelial recombination in *Rosa26-mTmG; Cdh5-CreERT2; Notch1^fl/fl^*mice and HV collection at P8. (B) Representative whole-mount confocal images of P8 HV following corn oil administration (CT; all cells are tdTomato⁺), 15 μg tamoxifen (mosaic recombination; GFP⁺/tdTomato⁺ cells), or 30 μg tamoxifen (complete recombination; GFP⁺ cells). (C) Bar graphs showing quantification of GFP⁺ and tdT⁺ vessel area as a percentage of the total vascular area (*n* = 3). (D) qRT-PCR analysis of *Notch1* mRNA expression in P8 HV following CT, 15 μg, or 30 μg tamoxifen treatment (*n* = 3). (E) Quantification of HV density (*n* = 3). (F) Representative confocal micrographs of P8 HV from *Rosa26-mT/mG; Cdh5-CreERT2* control mice following 15 μg of tamoxifen, showing GFP (green arrowheads), tdT (red arrowheads), and ephrin-B2 staining (n = 3). (G) Quantification of ephrin-B2⁺ cells among total tdT⁺ or GFP⁺ cells in *Rosa26-mT/mG; Cdh5-CreERT2* control HV (*n* = 3). (H) Representative confocal micrographs of P8 HV from *Rosa26-mTmG; Cdh5-CreERT2; Notch1^fl/fl^* mice following 15 μg of tamoxifen (*Notch1^cKO^*), showing GFP, tdT, and ephrin-B2 staining (*n* = 3). (I) Quantification of ephrin-B2⁺ cells among total tdT+ or GFP⁺ cells in *Notch1cKO* HV (*n* = 3). (J) Representative confocal micrographs of P8 HV from *Rosa26-mT/mG; Cdh5-CreERT2* control mice following 15 μg of tamoxifen, showing GFP, tdT, and EphB4 staining (*n* = 3). (K) Quantification of EphB4⁺ cells among total tdT⁺ or GFP⁺ cells in *Rosa26-mT/mG; Cdh5-CreERT2* control HV (*n* = 3). (L) Representative confocal micrographs of P8 HV from *Rosa26-mTmG; Cdh5-CreERT2; Notch1^fl/fl^* mice following 15 μg of tamoxifen (*Notch1^cKO^*), showing GFP, tdT, and EphB4 staining (*n* = 3). (M) Quantification of EphB4⁺ cells among total tdT⁺ or GFP+ cells in *Notch1^cKO^* HV (*n* = 3). (G, I, K, M) lines represent individual mice (*n* = 3 mice). Data are presented as mean ± SEM (*n* = 3 mice per group). Represented *p*-values are **≤ 0.01, ***≤ 0.001 and ****≤0.0001 from ordinary one-way ANOVA with Dunnett’s multiple comparisons test (C-E), or two-tailed parametric unpaired t-test (G, I, K, M). Non-significant (ns).

To verify the endothelial identity of GFP⁺ recombined cells, we stained P8 hyaloid vessels from tamoxifen-treated *Rosa26-mT/mG; Cdh5-CreERT2* control and *Rosa26-mT/mG; Cdh5-CreERT2; Notch1^fl/fl^* (*Notch1^cKO^*) mice for the endothelial nuclear marker ERG. Confocal analysis showed that most GFP⁺ cells expressed ERG in both genotypes (Fig. S4A, C), with approximately 90–95% of GFP⁺ cells being ERG⁺ (Fig. S4B, D), confirming the predominantly endothelial identity of the recombined population.

We next examined the relationship between the extent of endothelial recombination, Notch1 expression and hyaloid vessel persistence. Increasing the tamoxifen dose from 15 to 30 μg increased the proportion of GFP⁺ vascular area while reducing tdT⁺ vascular area (Fig. 6B, C). This was accompanied by reduced *Notch1* mRNA expression in isolated hyaloid vessels (Fig. 6D) and greater vascular persistence at P8 (Fig. 6E). Together, these findings establish an association between the extent of endothelial Notch1 recombination and the severity of the hyaloid regression defect.

To exclude a potential effect of the vehicle itself on endothelial arteriovenous identity, P8 *Rosa26-mT/mG; Cdh5-CreERT2; Notch1^fl/fl^*mice receiving corn oil alone were used as vehicle controls. Whole-mount immunostaining showed comparable ephrin-B2 and EphB4 expression patterns across the corn oil control conditions (Fig. S4E, F), indicating that corn oil administration did not alter the vascular distribution of these markers.

We next focused on the 15 μg mosaic condition to compare arteriovenous marker expression between GFP⁺ recombined and neighboring tdT⁺ unrecombined endothelial cells within the same vascular network. To determine whether Cre-mediated reporter recombination alone influenced ephrin-B2 expression, we first examined tamoxifen-treated *Rosa26-mT/mG; Cdh5-CreERT2* control mice carrying wild-type Notch1 alleles. Ephrin-B2 expression was detected in both tdT⁺ and GFP⁺ endothelial populations (Fig. 6F), with no significant difference in the percentage of ephrin-B2⁺ cells between the two populations (Fig. 6G). In contrast, in *Notch1^cKO^* mosaic vessels, ephrin-B2 was preferentially associated with tdT⁺ ECs and was reduced in GFP⁺ ECs (Fig. 6H). Accordingly, the percentage of ephrin-B2⁺ ECs was significantly lower in the GFP⁺ population than in the paired tdT⁺ population from the same mice (Fig. 6I). Similarly, EphB4 expression was examined in tamoxifen-treated Rosa26-mT/mG; Cdh5-CreERT2 control mice. EphB4 was detected in both tdT⁺ and GFP⁺ endothelial populations (Fig. 6J), with no significant difference in the percentage of EphB4⁺ cells between the two populations (Fig. 6K). Conversely, EphB4 was preferentially associated with GFP⁺ recombined ECs (Fig. 6L), with a significantly higher percentage of EphB4⁺ ECs in the GFP⁺ population than in the paired tdT⁺ population (Fig. 6M).

Because recombined and unrecombined endothelial cells were analyzed within the same vascular environment and no significant differences in ephrin-B2 or EphB4 expression were observed between these populations in control mice carrying wild-type *Notch1* alleles, these findings support a cell-autonomous role for endothelial Notch1 in regulating ephrin-B2 and EphB4 expression. Notch1-recombined endothelial cells preferentially displayed an ephrin-B2-low/EphB4-high profile, whereas neighboring unrecombined cells more frequently retained ephrin-B2 expression. These data support regulation of marker-defined arteriovenous features by Notch1 but do not establish direct transcriptional regulation or conversion of ephrin-B2⁺ cells into EphB4⁺ cells.

## DISCUSSION

Here, we identify dynamic changes in endothelial arteriovenous-associated marker expression during postnatal hyaloid vessel regression. The hyaloid endothelium displayed a predominantly ephrin-B2-positive profile at birth, followed by a progressive reduction in ephrin-B2 expression and enrichment of EphB4-positive endothelial cells as regression proceeded. These marker-defined populations were associated with distinct cellular behaviors: residual ephrin-B2-positive endothelial cells increasingly displayed C-Casp3 labeling, whereas EphB4-positive endothelial cells showed increased proliferative activity. Endothelial deletion of *Notch1* accentuated the ephrin-B2-low/EphB4-high profile but resulted in persistent hyaloid vessels. Mosaic recombination further showed that GFP-positive recombined endothelial cells preferentially expressed EphB4, whereas neighboring unrecombined cells more frequently retained ephrin-B2 expression. Together, these findings reveal that endothelial arteriovenous-associated features remain dynamic during developmental vessel regression and support a cell-autonomous role for Notch1 in regulating ephrin-B2 and EphB4 expression.

The mouse hyaloid vasculature has generally been described as an arterial vascular network. Our findings refine this view by showing that its arterial-associated profile is temporally regulated rather than maintained uniformly throughout regression. Ephrin-B2 expression was prominent at birth but declined markedly by P8, whereas EphB4 expression followed the opposite trajectory. Additional arterial- and venous-associated genes displayed marker-specific temporal changes, indicating that regression is not accompanied by a synchronous switch of an entire arteriovenous program. Instead, the hyaloid endothelium progressively changes its relative expression of arterial- and venous-associated features. This interpretation is consistent with increasing evidence that endothelial arteriovenous identity remains plastic during vascular development and remodeling[33] and can be reshaped by Notch activity, VEGF signaling and local hemodynamic cues[6, 14, 18, 34].

The endothelial localization analyses are important because ephrin-B2 and EphB4 expression was not restricted to the vascular compartment. Both proteins were also detected in hyaloid-associated extravascular cells, including macrophages. The combination of ERG with vascular labeling therefore allowed the endothelial changes to be distinguished from tissue-level alterations. Nevertheless, ephrin-B2 and EphB4 should not be interpreted as strictly exclusive lineage markers. Recent work has shown that their relative expression and signaling activities contribute to endothelial sorting and arterial specification, while allowing overlapping or context-dependent expression[18]. Accordingly, our results define changes in marker-associated endothelial features rather than demonstrating a direct conversion of ephrin-B2-positive arterial cells into EphB4-positive venous cells. Establishing such a transition would require simultaneous cell-resolved detection of both proteins combined with lineage tracing.

Our findings further associate these marker-defined populations with different endothelial behaviors. During physiological regression, C-Casp3 labeling increased among the residual ephrin-B2-positive endothelial population, whereas Ki67 positivity increased among EphB4-positive endothelial cells. These observations suggest that the changing endothelial composition is accompanied by a redistribution of apoptotic and proliferative activity. However, they do not establish that ephrin-B2 directly promotes apoptosis or that EphB4 directly drives proliferation. Indeed, Eph/ephrin signaling can elicit context-dependent effects on endothelial survival, migration and proliferation. EphB4 signaling has been implicated in endothelial proliferation and vascular morphogenesis[35–37], whereas ephrin-B2 signaling can regulate vessel pruning through STAT1-JNK3-dependent pathways[38]. Notably, previous disruption of ephrin-B2 signaling increased endothelial cell death while impairing vascular pruning[38], emphasizing that ephrin-B2 abundance, phosphorylation and downstream signaling are biologically distinct. Our experiments measure protein expression rather than pathway activation and therefore identify cellular associations that will require functional testing.

The effects of endothelial *Notch1* deletion provide further insight into the relationship between arteriovenous-associated expression and vessel regression. Notch signaling is a central regulator of arterial specification and can promote ephrin-B2 expression while restricting venous-associated programs[39–43]. Consistent with this function, *Notch1* deletion reduced endothelial ephrin-B2 expression and increased EphB4 expression in persistent P8 hyaloid vessels. Expression of *Dll4* was also reduced, whereas *Aplnr* and *Tek* expression increased. These findings support broader alteration of the arteriovenous-associated expression profile, although the whole-tissue RNA measurements may include contributions from non-endothelial cells.

Importantly, the ephrin-B2-low/EphB4-high profile observed after *Notch1* deletion resembles and accentuates the direction of the changes occurring during normal late regression. Nevertheless, *Notch1*-deficient vessels fail to regress[31]. The shift in marker expression is therefore not sufficient to induce vessel elimination. Instead, Notch1 appears to be required to coordinate endothelial molecular features with the cellular behaviors necessary for regression[44]. In persistent *Notch1^cKO^*vessels, apoptotic labeling was reduced within the residual ephrin-B2-positive population, whereas proliferation was increased within the expanded EphB4-positive population. Thus, Notch1 loss alters both the distribution of arteriovenous-associated markers and the behaviors associated with the corresponding endothelial populations. This altered coordination may contribute to the persistence of a proliferative endothelial compartment, although the present experiments do not demonstrate that changes in ephrin-B2 or EphB4 activity cause the regression defect.

The mosaic experiments provide evidence that the effect of Notch1 on ephrin-B2 and EphB4 expression has a cell-autonomous component. Graded tamoxifen administration increased endothelial reporter recombination, reduced *Notch1* expression and was associated with progressively greater hyaloid vessel persistence. Within the same mosaic vascular environment, GFP-positive recombined endothelial cells preferentially displayed an ephrin-B2-low/EphB4-high profile, whereas neighboring tdTomato-positive cells more frequently retained ephrin-B2 expression. Comparison of these populations within the same vessels minimizes differences in developmental stage, blood flow and local tissue environment. Nevertheless, recombination of the *Rosa26-mT/mG* reporter does not by itself confirm complete biallelic deletion of *Notch1* in every GFP-positive cell. Cell-resolved validation of Notch1 loss would further strengthen this conclusion. Moreover, whether Notch1 regulates *Efnb2* and *Ephb4* directly or through secondary transcriptional and biomechanical programs remains to be determined.

These findings may also be relevant to persistent fetal vasculature, in which failure of hyaloid vessel regression causes a spectrum of developmental ocular abnormalities[26, 27]. Our results do not establish that altered Notch1-ephrin-B2/EphB4 signaling causes human persistent fetal vasculature. They nevertheless identify an endothelial expression program that is altered when regression fails in mice and provide a framework for investigating whether similar changes occur in other experimental models or human PFV tissue. The observation that an ephrin-B2-low/EphB4-high profile can coexist with vascular persistence further suggests that the pathological outcome depends not simply on arteriovenous-associated marker expression, but on how these molecular features are coupled to endothelial survival, proliferation and multicellular remodeling.

Several limitations should be considered. First, ephrin-B2 and EphB4 expression defines marker-associated endothelial states but does not establish stable arterial or venous lineage identity. Second, because the two proteins were predominantly quantified in separate staining experiments, the abundance and behavior of double-positive or double-negative endothelial populations remain unresolved. Third, the apoptotic and proliferative analyses establish associations within marker-positive populations but do not demonstrate that Eph/ephrin signaling determines these behaviors. Direct comparison with marker-negative endothelial cells and functional perturbation of ephrin-B2 or EphB4 would address this question. Finally, rescue experiments will be required to determine whether restoring the ephrin-B2/EphB4 profile or modifying Eph/ephrin signaling can improve regression in *Notch1*-deficient vessels.

In conclusion, our study shows that hyaloid vessel regression is accompanied by dynamic remodeling of endothelial arteriovenous-associated features. The transition from an ephrin-B2-enriched to an EphB4-enriched endothelial profile is associated with changes in apoptotic and proliferative activity, while endothelial Notch1 is required to coordinate these molecular and cellular features during regression. By revealing that *Notch1*-deficient endothelial cells preferentially acquire an ephrin-B2-low/EphB4-high profile within mosaic vessels, these findings extend the role of Notch1 beyond arterial specification and identify a cell-autonomous contribution to endothelial heterogeneity during developmental vascular remodeling.

## MATERIALS AND METHODS

### Mouse strains

All animal procedures performed in this study were approved by the Institutional Animal Protection Committee (CIPA) of the Université du Québec à Montréal (UQAM), in accordance with the guidelines of the Canadian Council on Animal Care (CCAC). For inducible endothelial lineage tracing, Rosa26-mT/mG mice were bred with Cdh5-CreERT2 mice to generate *Rosa26-mT/mG; Cdh5-CreERT2* offspring. To achieve inducible endothelial-specific deletion of Notch1, *Notch1^fl/fl^*mice were crossed with *Cdh5-CreERT2* mice, generating *Notch1^fl/fl^; Cdh5-CreERT2* animals. *Notch1^fl/fl^* mice were kindly provided by Przemyslaw Sapieha and Nathalie Labrecque, whereas the *Cdh5-CreERT2* and *Rosa26-mT/mG* mouse lines were provided by Bruno Larrivée. All transgenic mouse lines were maintained on a C57BL/6J genetic background. Genotypes were determined by polymerase chain reaction (PCR) amplification of genomic DNA isolated from ear punchs, and animals carrying the appropriate alleles were selected for subsequent experiments.

C57BL/6J wild-type (WT) mice were obtained from The Jackson Laboratory.

### Tamoxifen injection

To induce Cre recombinase activity, tamoxifen (Sigma-Aldrich, Cat. No. T5648) was dissolved in corn oil (Sigma-Aldrich, C8267). For complete recombination, tamoxifen was prepared at 1 mg/mL, and pups received 30 µL per injection, corresponding to 30 µg of tamoxifen per injection. For mosaic recombination, tamoxifen was prepared at 0.5 mg/mL, and pups received 30 µL per injection, corresponding to 15 µg of tamoxifen per injection. In both experimental settings, tamoxifen was administered by intragastric injection into the milk spot once daily from postnatal day 1 (P1) to P3. For endothelial-specific *Notch1* deletion experiments, *Notch1^fl/fl^; Cdh5-CreERT2* pups received 30 µg of tamoxifen per injection and are referred to as *Notch1^cKO^*, whereas tamoxifen-treated Cre-negative littermates served as controls (*Notch1^cWT^*). For mosaic lineage-tracing experiments, *Rosa26-mT/mG; Cdh5-CreERT2* pups received 15 µg of tamoxifen per injection to induce partial Cre-mediated recombination and reporter switching from membrane-targeted tdTomato (mT) to membrane-targeted GFP (mG). Corn oil-treated *Rosa26-mT/mG; Cdh5-CreERT2* littermates were used as non-induced controls.

### Collection of pre and postnatal eyes for immunofluorescence

Following detection of the vaginal plug, pregnancy progression was monitored to determine the embryonic stage. Pregnant females were euthanized at embryonic day 18.5 (E18.5), and the uterine horns were exposed following a midline abdominal incision. Fetuses were carefully collected and removed from the amniotic sacs. E18.5 embryos, as well as pups at postnatal days 0 (P0), 4 (P4), and 8 (P8), were euthanized by decapitation. Eyes were subsequently enucleated, and the cornea was punctured with a 26G needle to facilitate fixation. The eyes were fixed in 4% paraformaldehyde (PFA) for 1 h at room temperature and washed three times with phosphate-buffered saline (PBS). Samples were either processed immediately or stored in PBS at 4 °C until further processing.

### Immunofluorescence

For immunofluorescence analysis of the hyaloid and retinal vasculature in embryonic and postnatal mice, eyes were enucleated and fixed in 4% PFA at room temperature, followed by three washes in cold 1× phosphate-buffered saline (PBS). Hyaloid tissues were permeabilized and blocked in PBS containing 10% FBS, 1% Triton X-100, and 0.01 M glycine. Retinas were incubated in a permeabilization and blocking solution containing 3% BSA, 3% FBS, 0.01% sodium deoxycholate, and 0.5% Triton X-100 in PBS. For blood vessel staining, hyaloids and retinas were incubated with isolectin B4 (IB4; 1:200; DL-1208.5, Newark, CA, USA) diluted in their respective blocking solutions. For protein localization, tissues were incubated with the appropriate primary antibodies (see Table S1), followed by the corresponding secondary antibodies: donkey anti-rabbit Alexa Fluor 488 IgG (1:500; A21206; Invitrogen), donkey anti-rabbit Alexa Fluor 594 IgG (1:500; A21207; Invitrogen), or donkey anti-rabbit Alexa Fluor 647 IgG (1:500; A31573; Invitrogen). Following immunostaining, tissues were washed with PBS and counterstained with DAPI (1 µg/mL; D9542; Sigma-Aldrich).

Retinas were directly flat-mounted, whereas hyaloid tissues were embedded in 2% ultra-low-melting-point agarose (800-016-TC) and incubated at 37 °C in a humidified chamber to allow gradual diffusion of the agarose. Hyaloids were subsequently transferred to 4 °C before flat-mounting. All samples were mounted in glycerol and imaged using an Olympus FV1000 confocal microscope with 10× and 60× oil-immersion objectives. Immunofluorescence quantification was performed using ImageJ on 3-5 Z-stack images of hyaloid vessels acquired at 60× magnification. For each animal, three confocal fields were quantified and averaged to obtain a single value per animal. Each data point therefore represents one independent animal. Cells were classified as extravascular based on their localization outside the IB4-positive vascular compartment, with no direct spatial contact with the IB4-positive vessel wall, as assessed throughout the confocal Z-stack and confirmed by 3D reconstruction. Images were processed using Adobe Photoshop 2020 (Adobe Inc).

### RNA extraction and real-time quantitative PCR

Total RNA was isolated from whole hyaloid vessels using the RNeasy Mini Kit (Qiagen, Germany) and eluted in RNase-free water. Following DNase I treatment, 1 µg of total RNA was reverse transcribed into complementary DNA (cDNA) using the SuperScript II Reverse Transcriptase kit (Thermo Fisher Scientific), according to the manufacturer’s instructions. For quantitative real-time PCR (qRT-PCR), 5 ng of cDNA was amplified in triplicate using either SYBR Select Master Mix or PowerUp SYBR Master Mix (Thermo Fisher Scientific).

Relative gene expression was determined using the comparative cycle threshold (ΔCt) method and normalized to the reference gene β-actin. Results are expressed as mean fold change ± SEM relative to the corresponding control samples. The sequences of the mouse-specific forward and reverse primers used in this study are provided in Table S2.

### Western blot

Hyaloid vessels were isolated from mice at embryonic day 18.5 (E18.5), and postnatal days 0 (P0), 4 (P4), and 8 (P8) for protein analysis. Samples from three mice were pooled to prepare each protein lysate. Tissues were mechanically disrupted in RIPA lysis buffer, and total protein concentrations were determined prior to electrophoresis. Equal amounts of protein were loaded onto polyacrylamide gels and separated by sodium dodecyl sulfate-polyacrylamide gel electrophoresis (SDS-PAGE).

Proteins were then transferred onto nitrocellulose membranes and blocked with 3% skim milk or 5% bovine serum albumin (BSA) in Tris-buffered saline containing Tween-20 (TBS-T) for 1 h at room temperature. Membranes were incubated overnight at 4 °C with the appropriate primary antibodies (Table S3), followed by incubation with the corresponding HRP-conjugated secondary antibodies for 1 h at room temperature. Protein signals were detected using Clarity ECL substrate (Bio-Rad) and visualized with a chemiluminescence imaging system.

Ephrin-B2 and EphB4 were analyzed on separate membranes. For each developmental stage, each independent biological preparation consisted of six HV pooled from 6 mice. Western blot experiments were performed in two to three independent biological preparations.

### Statistical analysis

Statistical analyses were performed using GraphPad Prism 8.0.2 (GraphPad Software, San Diego, CA). Data are presented as mean ± SEM or as individual values, as indicated. Prior to statistical testing, normal data distribution was assessed using the Shapiro-Wilk normality test with a significance level of 0.05. For comparisons between two groups, two-tailed parametric unpaired t-tests were used in a normal distributed data; otherwise, non-parametric Mann-Whitney U tests were applied. For comparisons involving more than two groups, ordinary one-way ANOVA followed by Dunnett’s multiple comparison test was used for normally distributed data. When data distribution did not follow, the non-parametric Kruskal-Wallis test followed by Dunn’s multiple-com parison correction was applied. It was considered statistically significant if the p values are *≤0.05, **≤0.01, ***≤0.001, ****≤0.0001. N is indicated pertinently for each experiment.

## SUPPLEMENTARY TABLES

**Table S1:** Primary antibodies used for Immunofluorescence.

| **<u>Antibody</u>** | **<u>Species</u>** | **<u>Clone</u>** | **<u>Dilution</u>** | **<u>Company</u>** | **<u>Catalog #</u>** |
| --- | --- | --- | --- | --- | --- |
| EphrinB2 | Rabbit | polyclonal | 1:200 | ABclonal | A5669 |
| EphrinB2 | Mouse | monoclonal | 1:200 | Santa Cruz Biotechnology | sc-398735 |
| EphrinB2 | Goat | polyclonal | 1:40 | R&D Systems | AF496 |
| EphB4 | Rabbit | polyclonal | 1:200 | ABclonal | A3293 |
| EphB4 | Mouse | monoclonal | 1:200 | Santa Cruz Biotechnology | sc-365510 |
| Ki67 | Rabbit | monoclonal | 1:300 | Cell Signaling | 9129S |
| C-CASP3 | Rabbit | monoclonal | 1:300 | Cell Signaling | 9664S |
| F4/80 | Mouse | monoclonal | 1:300 | Invitrogen | MF48000 |
| ERG1/2/3 | Rabbit | monoclonal | 1:300 | Abcam | ERP3864 |
| Anti-CD31-APC | Rat | monoclonal | 1:80 | BioLegend | 102510 |

**Table S2:**
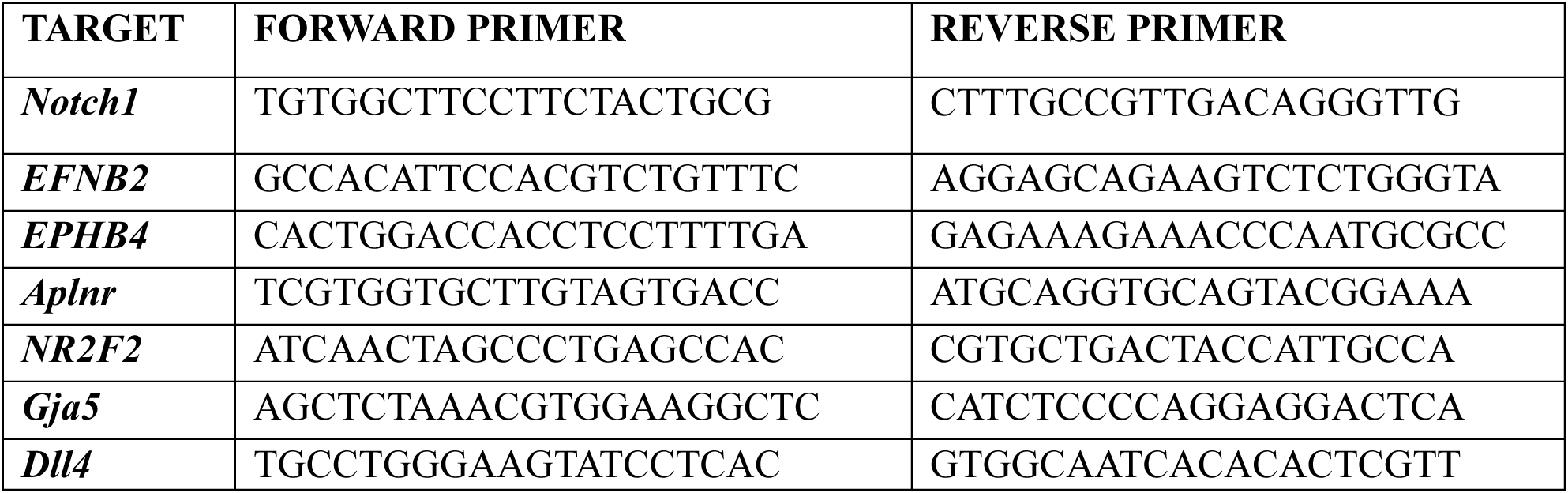
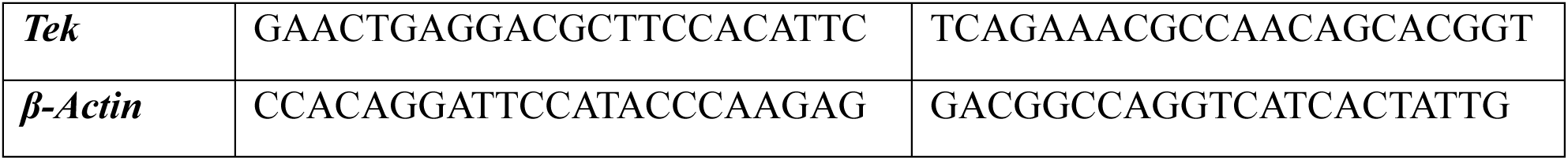
Table of oligonucleotides used for qRT-PCR reagents of murine origin.

**Table S3.** Primary antibodies used for Western blot analysis.

| **Antibody** | **Species** | **Clone** | **Dilution** | **Company** | **Catalog #** |
| --- | --- | --- | --- | --- | --- |
| β-actin | Mouse | monoclonal | 1:10000 | ABclonal | AC004 |
| NOTCH1 | Rabbit | polyclonal | 1:1000 | Cell Signaling | 3608 |
| EphrinB2 | Mouse | monoclonal | 1:1000 | Santa Cruz<br>Biotechnology | Sc-398735 |
| EphB4 | Mouse | monoclonal | 1:1000 | Santa Cruz<br>Biotechnology | Sc-365510 |

---

**Figure S1.**
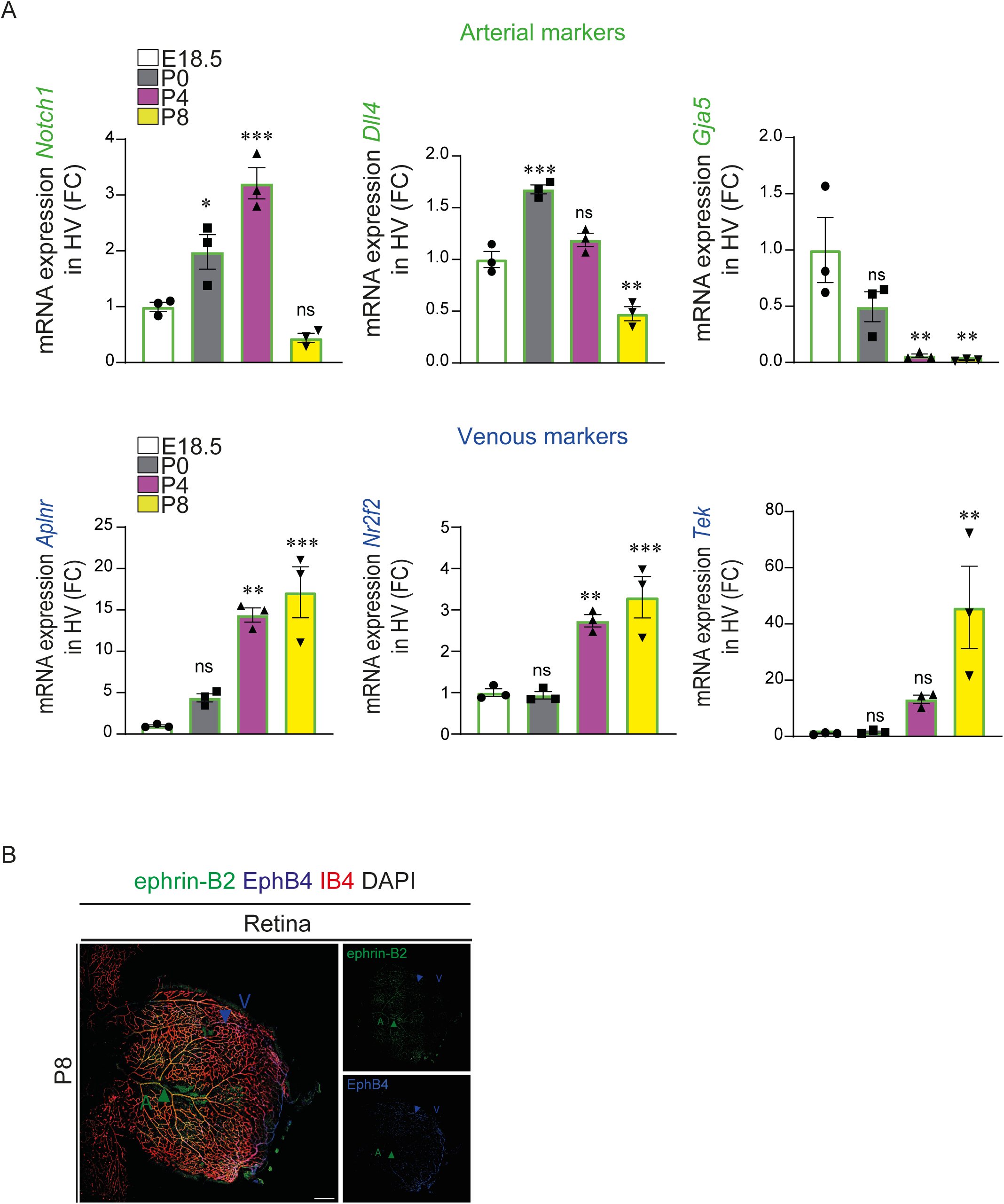
Characterization of arterial and venous marker expression during HV regression. (A) Quantitative RT-PCR analysis of HV collected at E18.5, P0, P4 and P8 showing the expression of and the arterial markers *Notch1*, *Dll4* and *Gja5,* and the venous markers *Aplnr*, *Nr2f2* and *Tek (n* = 3). (B) Representative confocal images of P8 retina labeled for ephrin-B2 (green), EphB4 (blue), IB4 (red), and DAPI (white), with retinal arterial (A) and venous (V) vessels shown as references for ephrin-B2 and EphB4 expression. *n* = 3 for *Notch1*, *Dll4*, *Gja5, Aplnr*, *Nr2f2* and *Tek* (A), with each point representing one independent biological preparation consisting of six HV pooled from three mice. Statistical significance was determined by ordinary one-way ANOVA followed by Dunnett’s multiple-comparisons test (A), with E18.5 as the reference group. *P* values are indicated as *≤ 0.05, **≤ 0.01, and ***≤ 0.001. Non-significant (ns). Scale bars, 500 μm (B).

**Figure S2.**
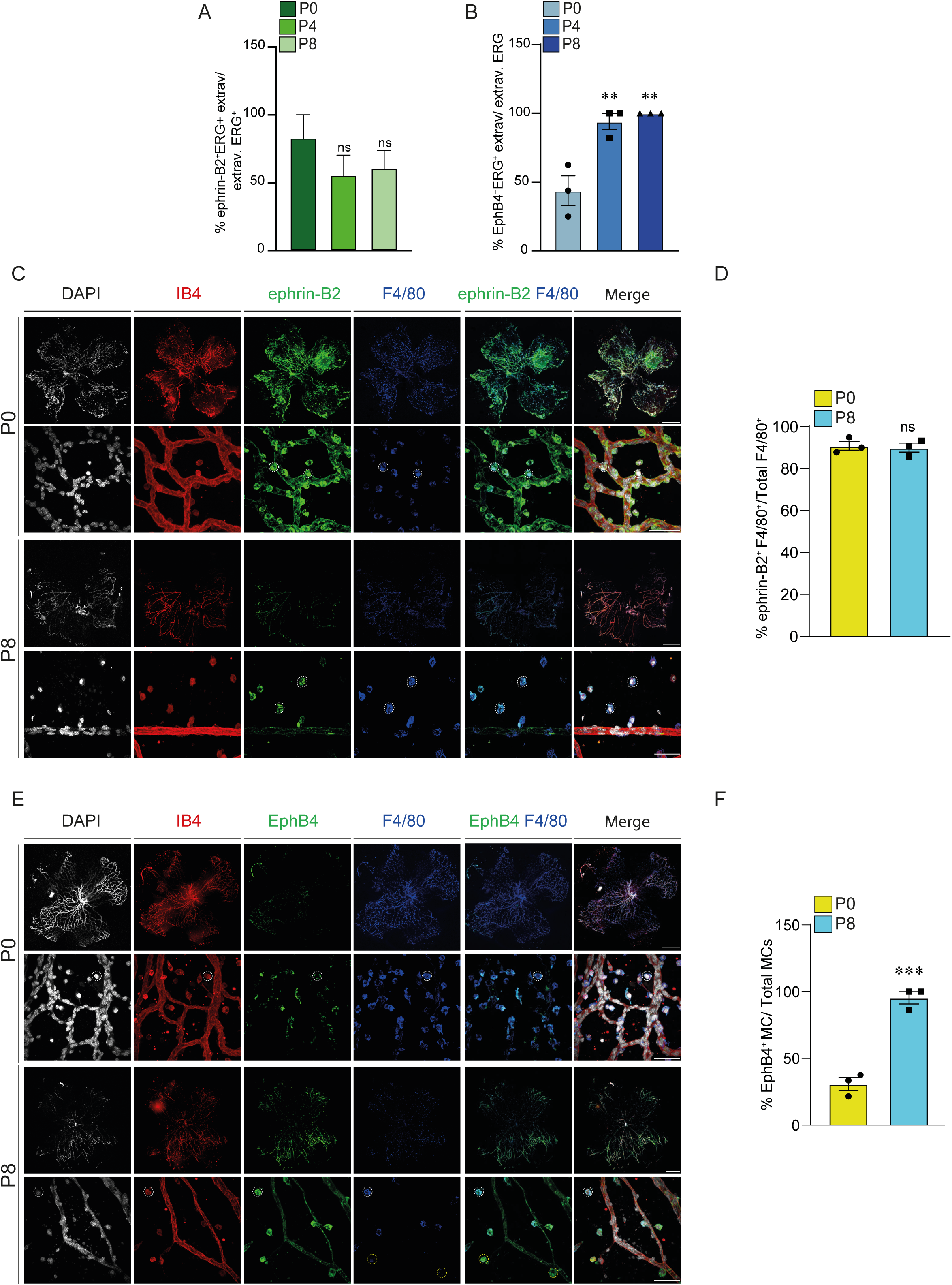
Expression of ephrin-B2 and EphB4 in extravascular cells and macrophages during hyaloid vessel regression. (A) Quantification of the proportion of extravascular ERG1/2/3⁺ephrin-B2⁺ cells and (B) extravascular ERG1/2/3⁺EphB4^+^ cells among total extravascular ERG1/2/3⁺ cells at P0, P4 and P8. (C) Representative confocal micrographs of whole-mount HV co-immunostained for IB4 (red), ephrin-B2 (green), F4/80 (blue) and DAPI (white) at P0 and P8 showing ephrin-B2 expression in F4/80⁺ cells. White circles indicate ephrin-B2⁺F4/80⁺ cells. (D) Quantification of ephrinB2⁺F4/80⁺ cells expressed relative to the total number of F4/80^+^ cells (n = 2-3). (E) Representative confocal micrographs of whole-mount HV co-immunostained for IB4 (red), EphB4 (green), F4/80 (blue) and DAPI (white) at P0 and P8 showing EphB4 expression in F4/80⁺ cells. White circles indicate EphB4⁺F4/80⁺ cells, and yellow circles indicate EphB4^+^F4/80^−^ cells. (F) Quantification of EphB4⁺F4/80⁺ cells among total number of F4/80^+^ (n = 2-3). Data are presented as means ± SEM. Represented *p*-values are **≤ 0.01 and ***≤ 0.001 from one-way ANOVA test with Dunnett’s multiple comparison (A, B), or two-tailed parametric unpaired t-test (D, F). Scale bars, 500 μm and 50µm [for higher-magnification images in (C, E)].

**Figure S3.**
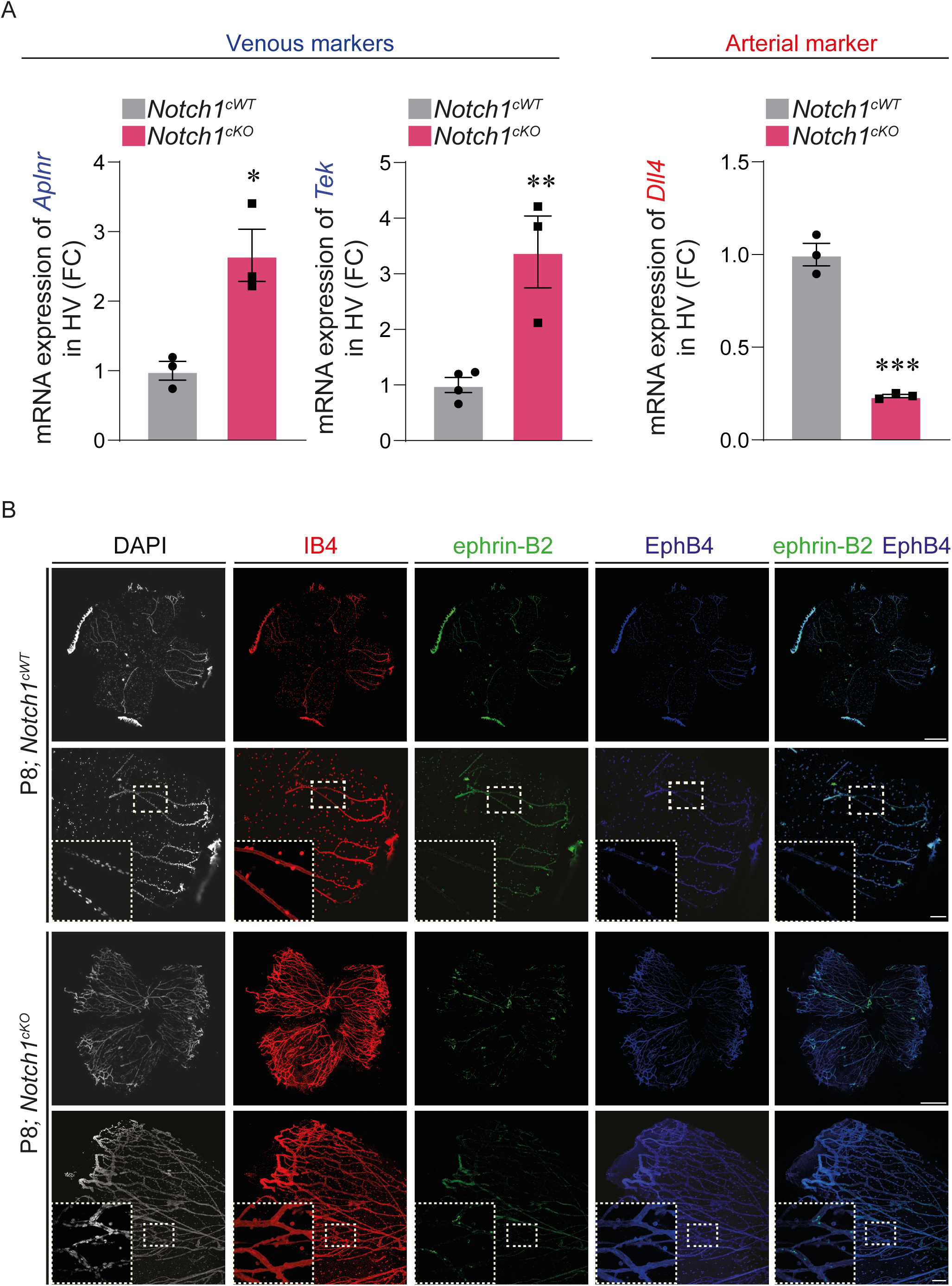
Endothelial Notch1 deletion is associated with increased venous-marker and reduced Dll4 expression in persistent hyaloid vessels. (A) Quantitative RT-PCR analysis of HV isolated from *Notch1^cWT^ and Notch1^cKO^*mice at P8 showing increased expression of the venous markers *Aplnr*, *Tek* and decreased expression of the arterial marker *Dll4* following endothelial-specific *Notch1* deletion (*n* = 3). (B) Representative images of ephrin-B2 (green), EphB4 (blue), IB4 (red) and DAPI (white) immunofluorescence staining of P8 HV *Notch1^cWT^* and *Notch1^cKO^* flatmounts (*n* = 4). Data are shown as means ± SEM. Represented *p* values are *≤ 0.05, **≤ 0.01, and ***≤ 0.001 from two-tailed parametric unpaired t-test. Non-significant (ns). Scale bars, 500 μm and 50 μm [for higher magnification images in (B)].

**Figure S4.**
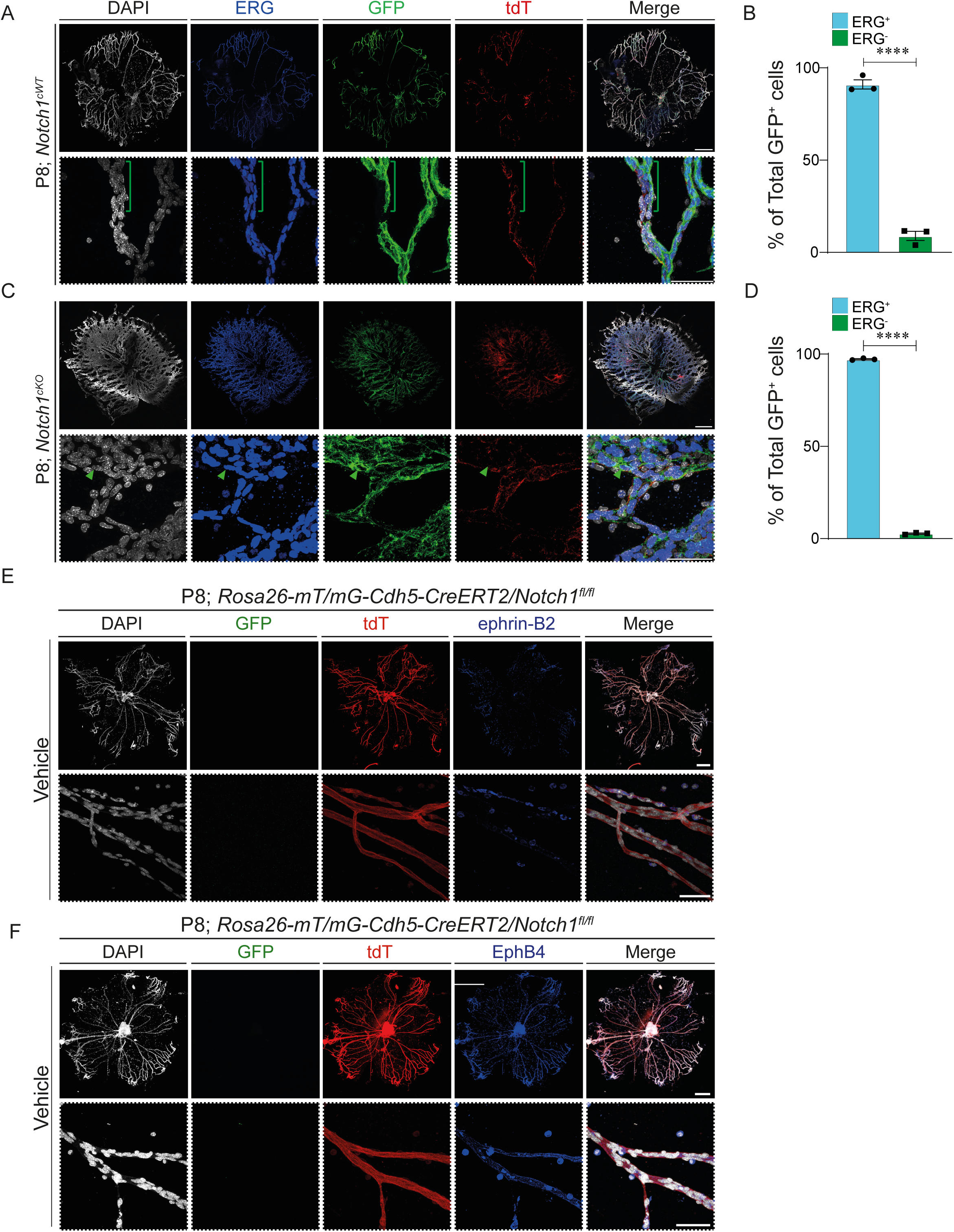
Endothelial Cre-mediated recombination doesn’t not alter ephrin-B2 and EphB4 expression in P8 hyaloid vessels. (A) Representative confocal images of P8 HV from *Rosa26-mT/mG; Cdh5-CreERT2* control mice following 15 μg of tamoxifen, labeled for ERG (blue), GFP (green), tdTomato (tdT; red), and DAPI (white) (*n* = 2). (B) Quantification of ERG⁺ and ERG⁻ cells among total GFP⁺ cells in tamoxifen-treated *Rosa26-mT/mG; Cdh5-CreERT2* control HV. (C) Representative confocal images of P8 HV from *Rosa26-mT/mG; Cdh5-CreERT2; Notch1^fl/fl^* mice following 15 μg of tamoxifen (*Notch1^cKO^*), labeled for ERG (blue), GFP (green), tdT (red), and DAPI (white) (*n* = 2). (D) Quantification of ERG⁺ and ERG⁻ cells among total GFP⁺ cells in tamoxifen-treated *Notch1^cKO^* HV. (E) Representative confocal images of P8 HV from *Rosa26-mT/mG; Cdh5-CreERT2/Notch1^fl/fl^* mice treated with 30 µL corn oil and labeled for GFP (green), tdT (red), ephrin-B2 (blue), and DAPI (white). (F) Representative confocal images of P8 HV from *Rosa26-mT/mG; Cdh5-CreERT2/Notch1^fl/fl^* mice treated with 30 µL corn oil and labeled for GFP (green), tdT (red), EphB4 (blue), and DAPI (white). Data are shown as means ± SEM. Represented *p* values are ****≤0.0001 from two-tailed parametric unpaired t-test. Scale bars, 500 μm and 50 μm [for higher magnification images in (A, C, E, F)].

## Notes

### Competing Interest Statement

The authors have declared no competing interest.

### Funding

- The authors acknowledge funding support from the Canada Research Chair/Canadian Institutes of Health Research (CRC-CIHR-00013897), the Natural Sciences and Engineering Research Council of Canada Discovery Grant (NSERC-DG-00000163), the Banting Foundation, the Centre d’Excellence en Recherche surles Maladies Orphelines-Fondation Courtois (CERMO-FC), and the Vision Health Research Network (VHRN).

